# *In silico* characterization of *GmSOS1* provides a comprehensive understanding for its role in soybean salt tolerance

**DOI:** 10.1101/2020.02.21.951061

**Authors:** Zhi-Chao Mei, Ling-Yan Yang, Zhi-Min Liu, Qi-Li Tang, Xin-Zhao Hou, Li-Jun Xie, Zhu-Jun Wei

## Abstract

Plant *SOS1* encodes plasma membrane Na^+^/H^+^ antiporter, which helps in the exclusion of Na^+^ and improves plant salt tolerance. However, detailed studies of *SOS1* in the important oil crop, soybean (*Glycine max*), are still lacking. In the present study, we carried out a comprehensive *in silico* analysis of *SOS1* in soybean. Referring to the analysis of physicochemical properties and structural characteristics, the GmSOS1 is an acidic protein with instability and hydrophobicity. Subcellular localization of *GmSOS1* supports the presumption that GmSOS1 is a plasma membrane Na^+^/H^+^ antiporter. Post-translational modification site prediction indicates 4 amino acids that may be phosphorylated. Further, the protein-protein interaction network and co-functional network signify the potential role of *GmSOS1* in salt stress tolerance. Although the interaction between *GmSOS1* and *GmHKT1* remains elusive, some of the intermediary signaling components of SOS pathway in soybean have been predicted. In addition, *in silico* expression analysis based on transcriptome datasets using publicly available database revealed that *GmSOS1* was differentially expressed in tissues and different times. Due to the analysis of its regulation mechanism, we found transcription factors such as WRKY and ERF as well as three miRNAs can regulate the expression of *GmSOS1*. Phylogenetic analysis using the homologous amino acid sequence of SOS1s from 26 species was performed to study the conserved motifs among these SOS1 members. Overall, we provide an extensive analysis of the *GmSOS1* and it promises the primary basis for the study in development and response to salt tolerance.

## 1. Introduction

High concentrations of Na^+^ in saline soils inhibit plant growth and reduce agricultural productivity^[1]^. Salt stress causes primary injuries, including osmotic effects at an early phase and ionic toxicity at a later phase of plant growth^[2]^. It can also induce the accumulation of reactive oxygen species (such as O^2-^ and H2O2), which will lead to secondary damages such as nutritional imbalance and oxidative stress^[3–5]^. Through long-term evolution, plants have developed diverse mechanisms to mitigate the destructive effects of salt stress through morphological, physiological and biochemical adjustments^[6–7]^. Sodium (Na^+^) is a common ion in soil, and excessive Na^+^ outside of or accumulation in plant cells results in disturbance or imbalance of intracellular osmotic, ionic, and oxidative homeostasis^[8]^. Thus, it is essential for plants to prevent the accumulation of Na^+^ and to maintain the appropriate K^+^/Na^+^ ratio in the cytoplasm, which is largely regulated via Na^+^ transporters^[19-12,2]^. SOS (Salt overly sensitive) pathway can specifically regulate cell sodium homeostasis, which is one of the most extensively studied plant salt stress signal transduction pathways^[13]^. Under high Na^+^ stress, calcium-dependent protein kinases in the SOS pathway mediate salt stress signals, and plants can adapt salt stress by effluxing Na^+^. The SOS pathway consisting of three components, the plasma membrane Na^+^/H^+^ antiporter SOS1, the cytoplasmic protein kinase SOS2, and the Ca^2+^ sensor SOS3. SOS1 protein plays an important role in SOS pathway to efflux Na^+[14]^. Up to now, a number of *SOS1* homologous genes have been cloned and identified from a variety of different plants. The *A. thaliana* SOS1 protein (AtSOS1) is the first described plasma membrane-localized Na^+^/H^+^ antiporter mediating Na^+^ efflux and controlling long-distance Na^+^ transport from roots to shoots^[15]^. When tomato (*Solanum lycopersicum) SlSOS1* was completely silenced, the plant showed a salt-sensitive phenotype with high Na^+^ content in roots and leaves. However, when *SlSOS1* was specifically silenced in tomato stem vascular bundles, the plant showed a salt tolerance phenotype with low Na^+^ content, indicating that the SlSOS1 protein mainly excreted Na^+^ in tomato root cells^[16]^.

Soybean is the primary source of the world’s supply of vegetable protein and oil. Demand for soybean has grown rapidly due to soybean’s wide range of applications in food, feed, and industrial products. However, soybean production is threatened by several abiotic stresses, such as soil salinity. Soybean *GmSOS1* can reduce primary Na^+^ toxicity by limiting Na^+^ accumulation and enhance antioxidant enzyme activity to reduce secondary oxidative stress^[17]^. In present study, a comprehensive bioinformatics analysis of soybean *GmSOS1* gene was performed, which lays the foundation for further research on the biological properties and functions of soy and provides a theoretical basis for the future breeding of high-yield, salt-tolerant soybean varieties.

## 2. Materials and methods

### 2.1 Genome-wide identification

The plant SOS1 protein has an N-terminal Na^+^/H^+^ exchanger domain (PF00999) and a long cytoplasmic tail at C-terminal end. The cytosolic moiety has the cyclic nucleotide-binding domain (PF00027) and the auto-inhibitory domain, which can regulate the transport activity. To identify the *SOS1* gene in soybean, genome and proteome datestes were downloaded from Soybase^[18]^, and the Hidden Markov Model (HMM) profiles of the first two abovementioned domains were downloaded from Pfam 32.0 datebase^[19]^. The two HMM profiles were used as query to search the soybean proteome with Hmmer 2.0(http://www.hmmer.org/).

### 2.2 Analysis of protein sequence properties

The protein sequence of *GmSOS1* gene was used in ExPASy tools^[20]^ with standard parameters to calculate the physicochemical properties. Subcellular localization was predicted by WoLF PSORT^[21]^, Plant-mPLoc^[22]^ and ProtComp9.0 (http://www.softberry.com/) independently.

### 2.3. Protein structure analysis

Raptorx-propert^[23]^ and Netsurfp-2.0^[24]^ were used to predict secondary structure, solvent accessibility and disordered regions of GmSOS1 protein. Typical domains were indentified by PfamScan^[25]^ and CDD^[26]^, the specification of the membrane spanning segments and their IN/OUT orientation relative to the membrane was predicted by TOPCONS^[27]^, and signal peptides was predicted by signalp-5.0 Server^[28]^. We aligned the protein sequences of GmSOS1 and AtSOS1 to deduce the location of auto-inhibitory domain in GmSOS1^[29]^. The three-dimensional (3D) structure of GmSOS1 was generated by RaptorX ^[30]^, and VMD^[31]^ was used to visualize the structure. The ligand-binding site of CNBD was predicted by 3DligandSite^[32]^.

### 2.4 Protein phosphorylation site analysis

In order to ensure the high reliability of protein phosphorylation sites prediction results, local prediction software GPS5.0^[33]^ and two online prediction services NetPhos 3.1Server^[34]^ and MusiteDeep^[35–36]^ were used independently for analysis. Among them, the confidence parameter of GPS5.0 was set as “high”, NetPhos 3.1 Server only displays the residues whose scores were higher than 0.90, and the lowest specificity of each predicted site discovered by MusiteDeep was set as “90%”. The sites indicated by all the three results were selected.

### 2.5 Protein function analysis

The functional protein association network of GmSOS1 protein was predicted using the web program STRING ^[37]^, the confidence parameter was set as “high confidence”. Genes co-functioned with *GmSOS1* gene were accessed by SoyNet^[38]^. Visualizing was performed with Cytoscape tool^[39]^.

### 2.6 Expression profiles and its regulation mechanism

#### 2.6.1 The spatio-temporal and stressed expression profiles of GmSOS1 gene

Publicly available soybean transcriptome datasets from Phytozome v12.1^[40]^ were used to investigate the expression pattern of *GmSOS1* gene in nine different tissues including roots, root hairs, stems, nodules, apex meristem, leaves, flowers, pods, and seeds. The circadian transcriptome of soybean leaves (GSE94228), salt stress transcriptome of soybean leaves (GSE133574) and short-term phosphorus deficiency of soybean leaves (GSE104286) also been collected from NCBI GEO^[41]^. The *GmSOS1* gene expression level was presented as FPKM (fragments per kilobase of exon per million fragments mapped). Two-tailed test was used for the comparison and *P* < 0.01 was considered as statistically significant.

#### 2.6.2 Transcription factor analysis and miRNA predicted

The upstream 2kb sequences of the *GmSOS1* from the translation start site were retrieved to analyze for the identification of transcription factors important for gene expression under abiotic stress using PlantPAN 3.0^[42]^. We used the sRNAanno^[43]^ to study the role of miRNAs in regulating the expression of *GmSOS1* gene.

### 2.7. Conserved motif and phylogenetic analysis

The homologous amino acid sequence of SOS1s from 26 species were derived from Phytozomev12.1^[40]^ and BLAST program of NCBI^[44]^. They were then aligned using the MEGA^[45]^-MUSCLE^[25]^program. Aferwards, a phylogenetic tree was constructed for the study of the these SOS1 members with the neighbor-joining(NJ) method. Bootstrap analysis was performed with 1000 replicates for statistical reliability. There were 26 species in the phylogenetic tree, comprising 15 SOS1s from Leguminosae (*Glycine max、 Glycine soja、 Vigna radiata、 Spatholobus suberectus、 Phaseolus vulgaris、 Vigna angularis、 Vigna unguiculata、 Abrus precatorius、 Cicer arietinum、 Pisum sativum、 Trifolium repens、 Melilotus officinalis、 Medicago truncatula、 Lupinus angustifolius、 Arachis hypogaea)*, 2 from Solanaceae (*Oryza sativa、 Brachypodium distachyon*), 1 from Cruciferae (*Arabidopsis thaliana*), 1 from Bromeliad (*Ananas comosus*), 2 from Gramineae (*Oryza sativa、 Brachypodium distachyon*),1 from Vitis(*Vitis vinifera*),1 from Rosaceae(*Prunus persica*), 1 from Myrtaceae (*Eucalyptus grandis*), 1 from Willow branch (*Populus trichocarpa*), and 1 from Mallow family(*Gossypium raimondii*). The motifs of these proteins were identified using the MEME^[46]^ program. The analysis was performed using “searching for six motifs” and the optimum motif width was set to 6–50 while the rest of the settings were kept as default.

## 3. Results and analysis

### 3.1 Genome-wide identification

There was only one gene encodes the protein which contained the Na^+^/H^+^ exchanger domain and cyclic nucleotide-binding domain, it was located on Chr8(+):6947483-6960485. The exon number of *GmSOS1* gene was 23.

### 3.2. Analysis of protein sequence properties

The results showed that the formula of this protein was C_5708_H_8995_N_1513_O_1650_S_38_, the molecular weight (Mw) was 12.64kDa. The theoretical isoelectric value (pI) was 6.30, which revealed that it was an acidic protein. The GmSOS1 polypeptide was an 1143 amino acid sequence composition bias toward leucine (12.69%). In addition, there was an apparent lack of tryptophan (1.40%) and cysteine (1.14%). The number of positively charged residues (Arg+Lys) was 104, while the number of negatively charged residues (Asp+Glu) was 117 (Figure 1). The instability index (II) of GmSOS1 was computed to be 42.61, this classified the protein as unstable. The grand average of hydropathicity (GRAVY) was 0.089, implying that it was hydrophilic protein. The Aliphatic index (AI) is 102.29, which also proves GmSOS1 is hydrophilic protein. All three subcellular localization prediction methods showed that GmSOS1 was localized at the plasma membrane, consistent with the presumption that GmSOS1 protein is a plasma membrane Na^+^/H^+^ antiporter.

**Figure 1.**
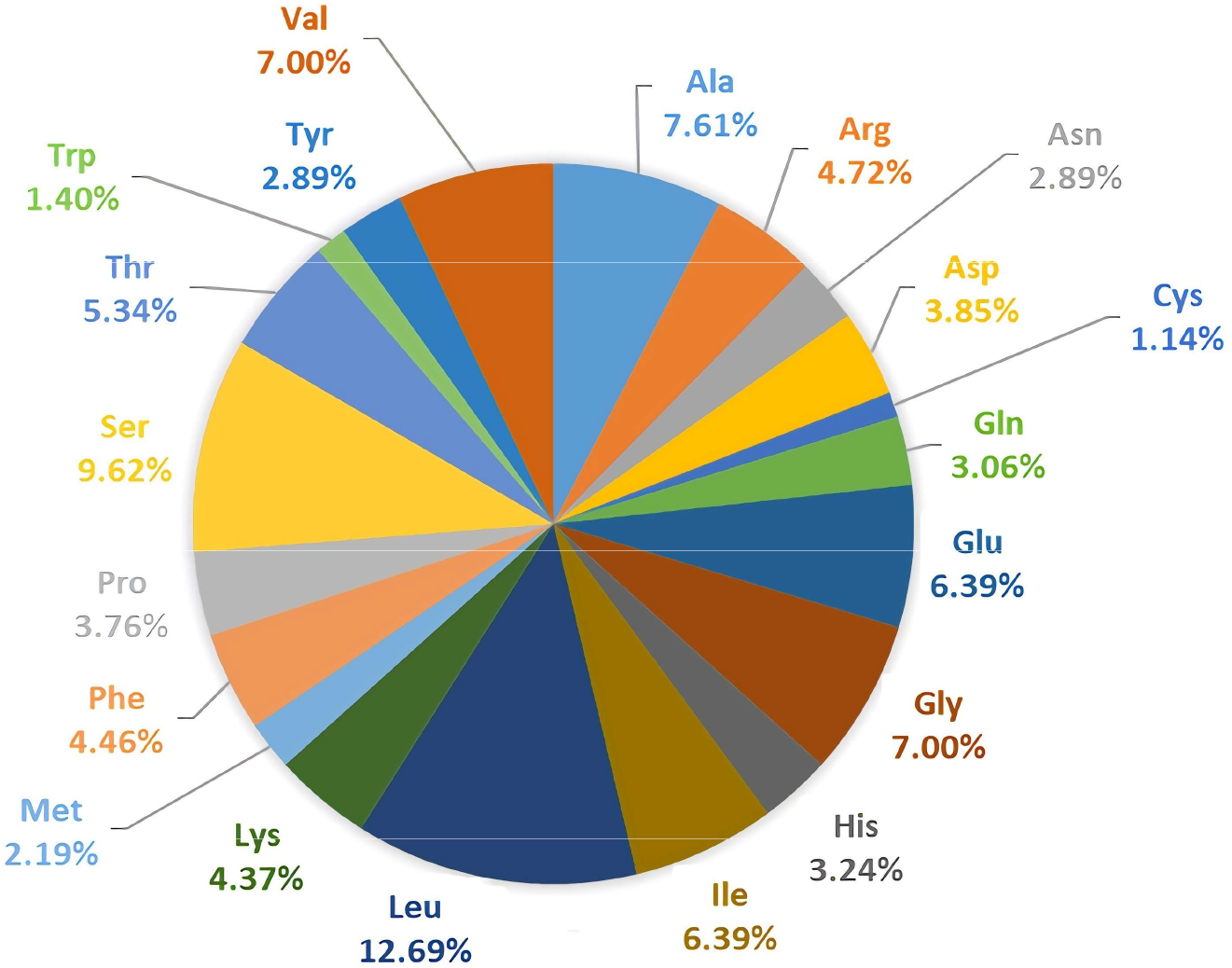
amino acid composition of GmSOS1 protein.

### 3.3 Protein structure analysis

The deduced secondary structure of GmSOS1 protein indicated that it formed of 54% helixes, 7% strands and 37% coils. The amino acids exposed to the protein surface and embedded in the protein accounted for 49% and 23% of the total amino acids, respectively. 20% of the amino acids constituted the disordered region of the protein structure. The N-terminal of the protein accounted for about two-thirds of the total peptides was mainly composed of helixes, the near-C-terminal in the middle occupied one-sixth of the total peptides and consisted of two strand regions and a helical region separating them. 1/6 positions in the C-teminal predicted as disordered with high solvent accessibility (Figure 2). The Na^+^/H^+^ exchanger domain was located at the N-terminal (32-464) which included 13 transmembrane regions (TM1:32-52, TM2:58-78, TM3:95-115, TM4:128-148, TM5:158-178, TM6:194-214, TM7:227-247, TM8:278-276, TM10:314-334, TM11:351-407, TM13:425-445). A cyclic nucleotide-binding domain was found near C-terminal (763-844). GmSOS1 protein also contained a C-terminal auto-inhibitory domain (994-1143). Signalp-5.0 Server showed no significant signal peptides (Figure 3A). The visualized 3D structure of the protein was shown (Figure 3B). The N-terminal of the Na^+^/H^+^ exchanger domain was a hydrophobic structure, and the C-terminal had a hydrophilic tail. The ligand CMP of the CNBD domain was bound to a hydrophobic pocket on the surface of the domain (Figure 3C).

**Figure 2.**
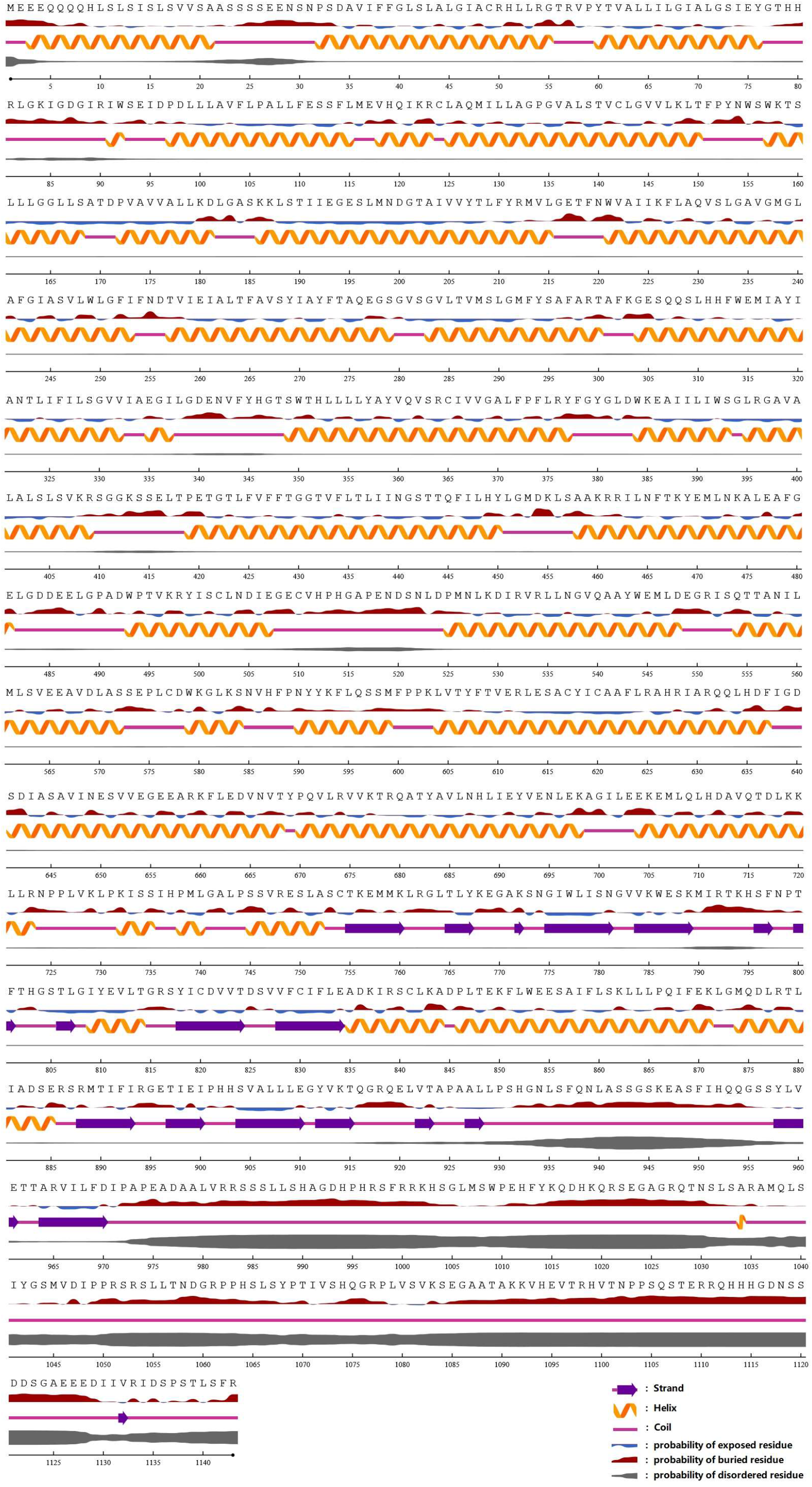
secondary structure, solvent accessibility and disorder regions of GmSOS1 protein.

**Figure 3.**
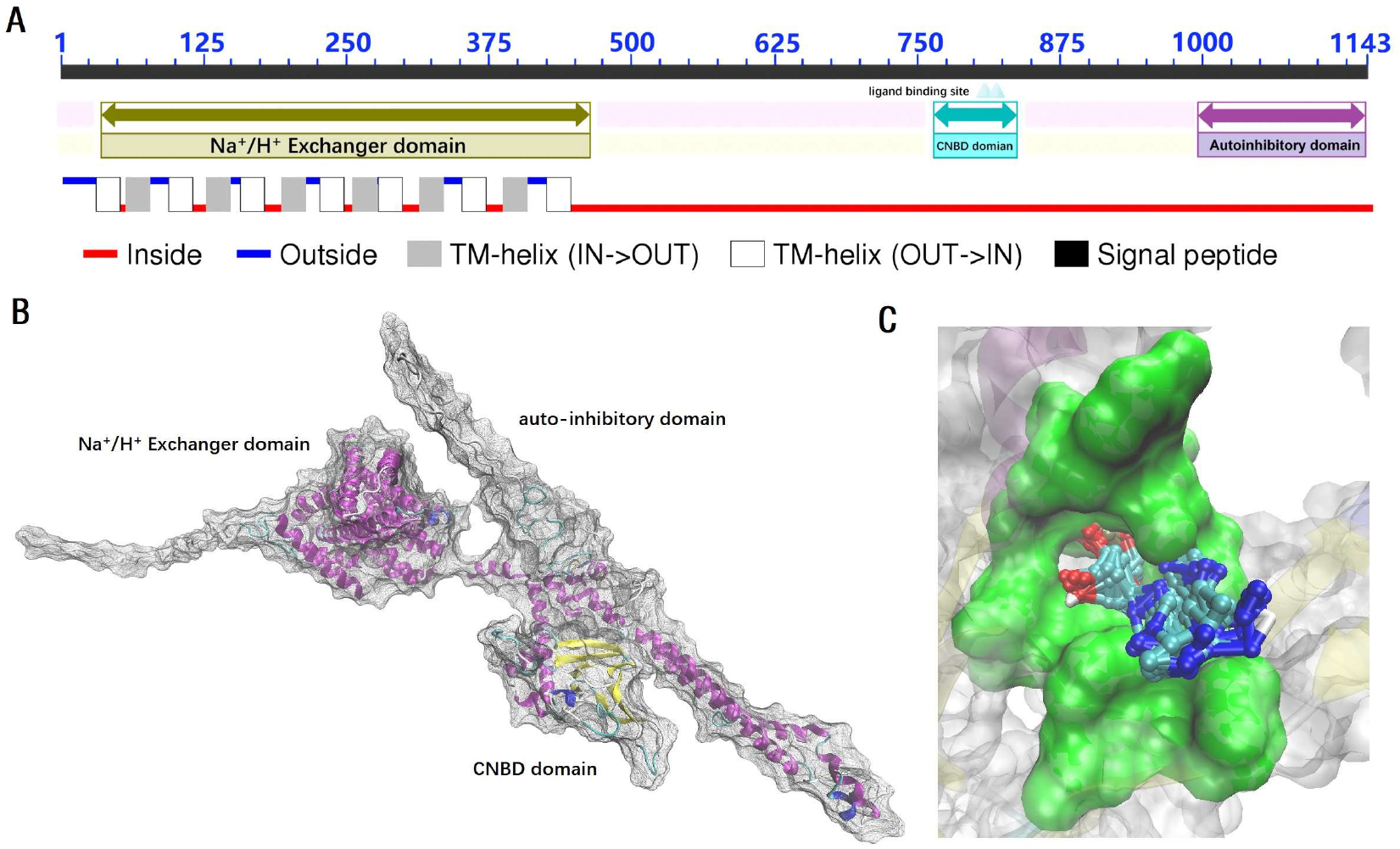
protein structure of GmSOS1.

### 3.4 Protein phosphorylation site analysis

All three methods for putative phosphorylation sites in GmSOS1 identified several consensus sequences at the C-terminal hydrophilic domain, and the amino acid positions were S24, S414, S985, S1119, S1123, and S1136, respectively. Further transmembrane domain analysis showed that only the last four serines residues are target phosphorylation sites in the cytoplasm which may play an important role in the regulation of GmSOS1 protein activity (Figure 4).

**Figure 4.**
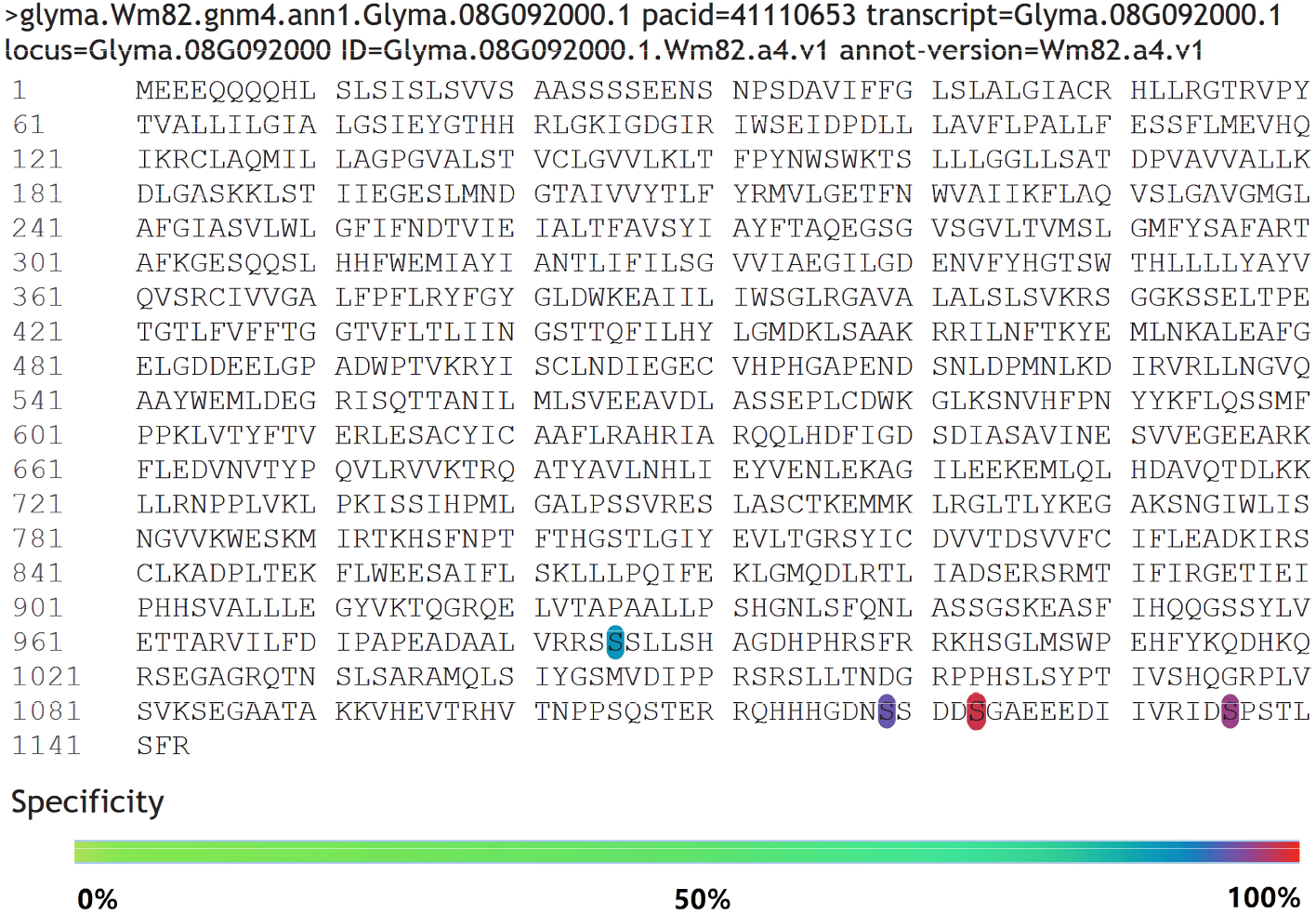
phosphorylation sites of GmSOS1 protein.

### 3.5 Protein function analysis

The functional protein association network was predicted in order to gain a better understanding of GmSOS1 protein role in soybean. The results showed that GmSOS1 protein was mapped to 10 unknown proteins that participated in the interaction network (Figure 5A). Comparative analysis of homologs between Arabidopsis and soybean may be helpful for the understanding of homologous gene functions in soybean. Our results showed that Glyma.01G002300.2, Glyma.06G271600.1’Glyma.07G130100.2,Glyma.12G133400.1 and Glyma.17G31011.1 (Wm82.a1.v1) might be function as *A. thaliana* AtHKT1 (High-affinity K^+^ channel transporter). The K^+^-Na^+^ co-transporter HKT1 also functions in Na^+^ influx under salt stress^[47]^, while SOS1 is a Na^+^-efflux transporter. At the same time, a high cytosolic K^+^/Na^+^ ratio is essential for maintaining cellular metabolism ^[48]^, and the regulation mechanism between SOS1 and HKT1 remains to be explored. Glyma.17G229500.1 and AtCBL10 shared the conserved EF-hand calcium binding domain. It is worthy to note that the RST and PARP domains were found in Glyma.01G015000.8 and Glyma.09G207200.4 as well as AtRCD1. Glyma.10G018000.1 had the Na^+^/Ca^2+^ exchanger domain. Glyma.01G015200.2 had no special domain. Co-functional networks are useful for identifying genes that are involved in a particular pathway or phenotype. To understand the interactions of *GmSOS1* genes with its neighbors and to get more insights into its function, a gene correlation network was constructed. It can be seen that the *GmSOS1* may be directly related to 12 genes (Figure 5B), nine of which are widely connected with other genes in gene co-functional network and regulated diverse biological processes and pathways. Notably, the co-functional network was divided into 4 clusters. In cluster I, Glyma.06G128700 is the homologous gene of *AtCBL4 (SOS3)* and Glyma.04G235900 is the homologous gene of *AtCBL7*. In cluster II, Glyma.17G113700 and Glyma.13G166100 are homologous genes of *AtCIPK24 (SOS2)* annotated in Soybase. Cellular calcium signals are detected and transmitted by sensor molecules such as calcium-binding proteins. In plants, the calcineurin B-like protein (CBL) family represents a unique group of calcium sensors and plays a key role in decoding calcium transients by specifically interacting with and regulating a family of protein kinases (CIPKs)^[49]^. Based on two proven salt tolerance pathways in Arabidopsis, one is the phosphorylation of Na^+^/H^+^ antiporter SOS1 on the plasma membrane by the *AtCBL4 (SOS3)-AtCIPK24 (SOS2)* kinase complex brings about substantial activation of SOS1 to enhance Na^+^ exclusion^[50]^, and the other is through *AtCBL10-AtCIPK24 (SOS2)* kinase complex phosphorylates Na^+^/H^+^ antiporter NHX on the vacuolar membrane^[51]^. In this study, *GmCBL4* has the same fun ction as *SOS3, GmCBL10* has the same function as *AtCBL10*, and a new *GmCBL7* with similar function as *SOS3* is predicted. Glyma.01G015000, Glyma.09G207200 and Glyma.01G15200 are in cluster 3, all of which are homologous genes of *AtRCD1*. In protein-protein interaction analysis, it can be seen that the proteins encoded by the first two genes interact with GmSOS1. It was found that the *AtRCD1* in Arabidopsis was mainly involved in plant regulation of salt stress through two pathways. One is a function in the nucleus. It has been shown that *RCD1* interacts with transcription factors such as STO and DREB2A, which have been implicated in salt-stress tolerance ^[52]^. Another function of RCD1 is performed in the cytoplasm and near the cell periphery and is possibly related to its interaction with SOS1, the biochemical consequence of RCD1–SOS1 interaction is not known^[53]^. Glyma.05G097300 encoded type D cyclin, which was localize to the nucleus. Glyma.08G219500 encoded glycosyltransferase. Glyma.08G165800 was a protein kinase, the function of Glyma.09G259200 and Glyma.06G027800 were unknown.

**Figure 5.**
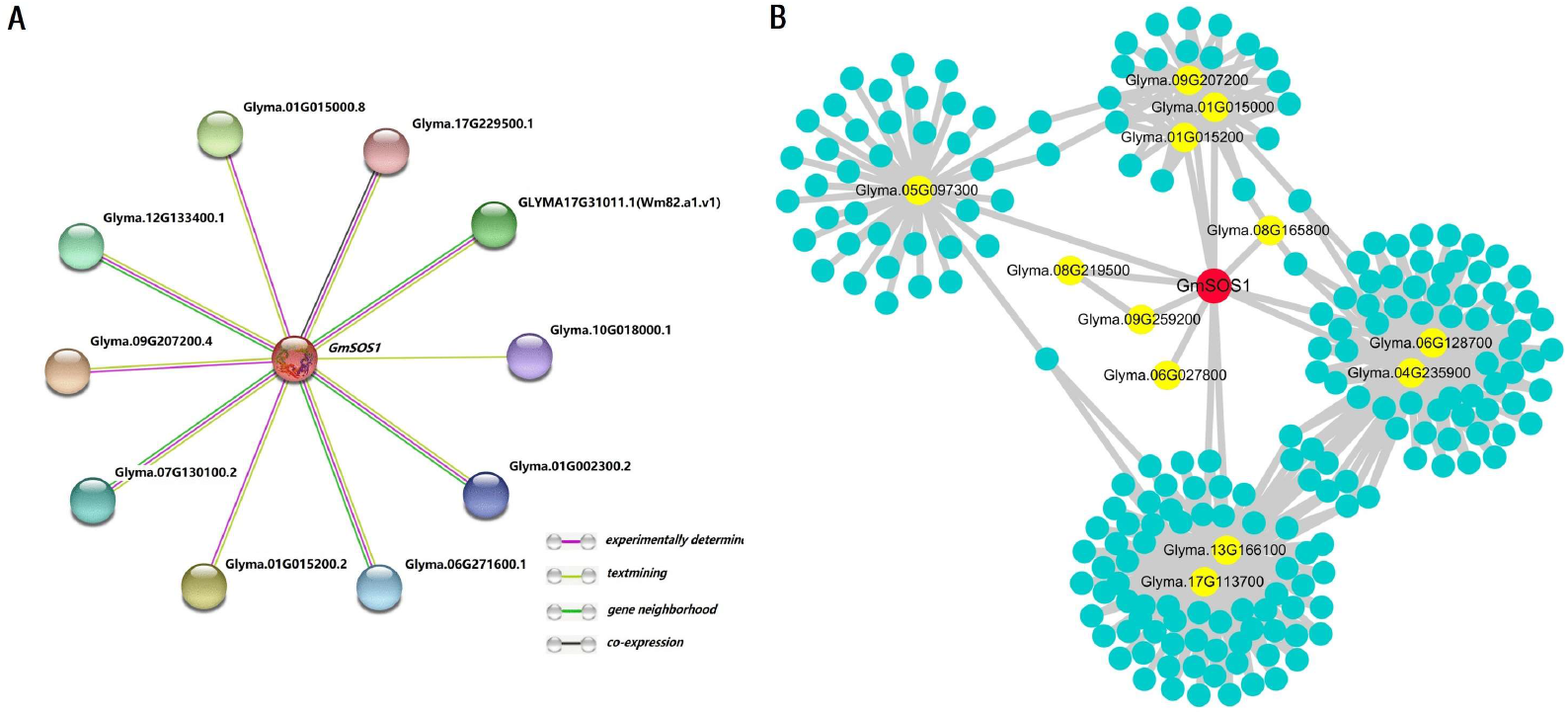
The functional protein association network and co-functional network of *GmSOS1*.

### 3.6 Expression profiles and its regulation mechanism

#### 3.6.1 The spatio-temporal and stressed expression profiles of GmSOS1 gene

*GmSOS1* showed the highest level of expression in roots and the lowest in leaves (Figure 6A). On the whole, the expression levels of *GmSOS1* varied at different time, suggesting that the expression of *GmSOS1* may have circadian rhythm (Figure 6B). *GmSOS1* was slightly up-regulated (*p* > 0.01) when treated with phosphorus deficiency in soybean leaves (Figure 6D). However, it was significantly induced in root (*p* < 0.01) under salt stress, which indicates its role involved in salt responses (Figure 6C).

**Figure 6.**
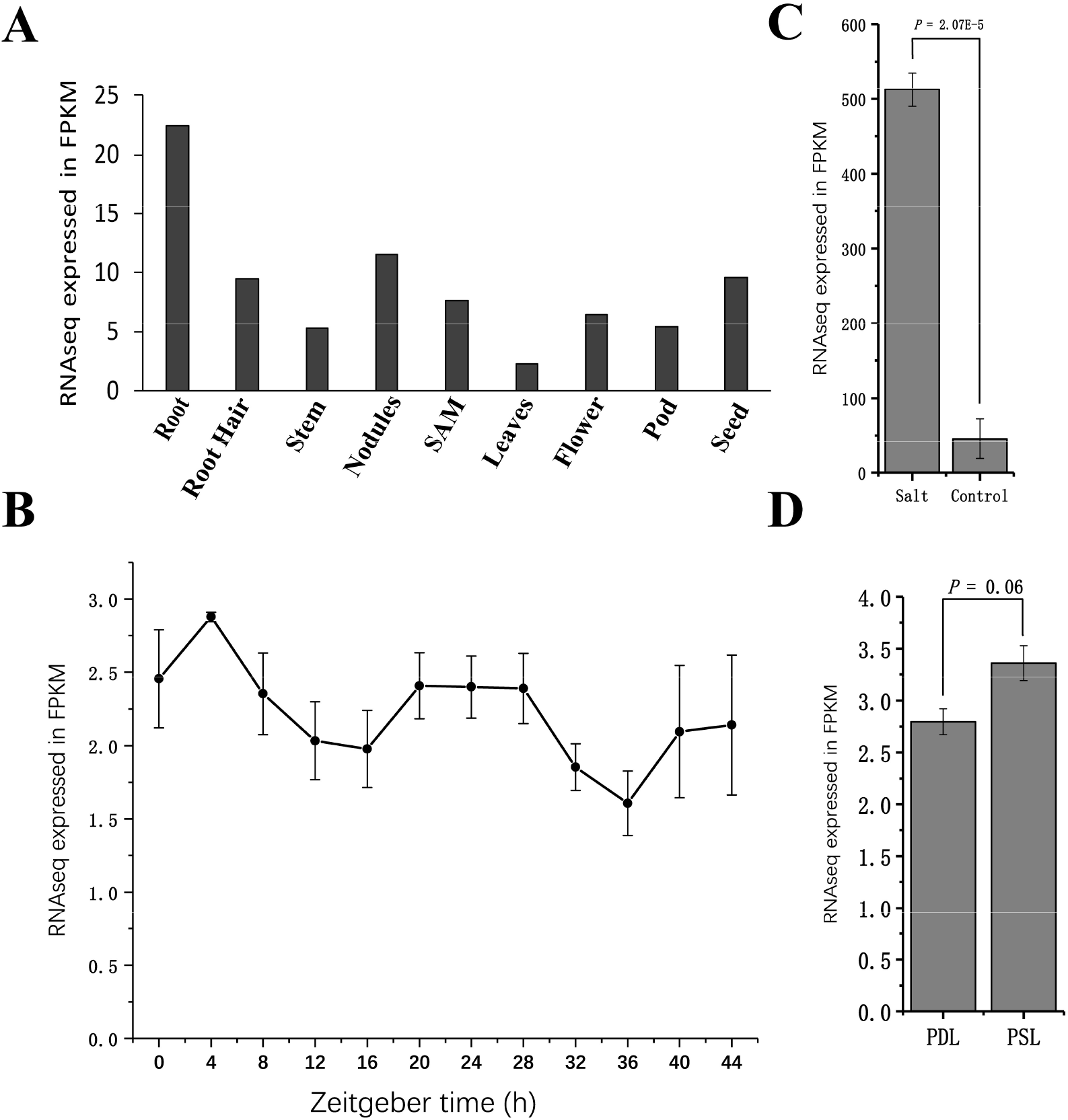
The spatio-temporal and stressed expression profiles of GmSOS1 gene. PDL: phosphorus deficiency in leaves PSL:phosphorus deficiency in leaves(means ± SDs)

#### 3.6.2. Transcription factor analysis and miRNA predicted

The results showed that 27 transcription factors that participated in the regulation of *GmSOS1*, including 2 ARR-B families, 1 bZIP family, 2 C2H2 families, 4 ERF families, 3 HD-zip families,1 MIKC-MADS family, 1 MYB-related family, 2 NAC families, 1 TBP family, and 11 WRKY families (Figure 7). *GmWRKY47* transcription factor was identified from the soybean genome via a genome-wide study, Yu et al.^[54]^ found that Glyma.09G080000 encoded by *GmWRKY47* was involved in the response to salt stress. In this study, we found that the *GmWRKY47* can probably target *GmSOS1*, which may be a potential mechanism for *GmWRKY47* to participate in salt stress response. We predicted *GmSOS1* gene as potential target of 3 miRNAs: Gma-mir1508, Gma-mir395, and Gma-mir166 (Figure 7).

**Figure 7.**
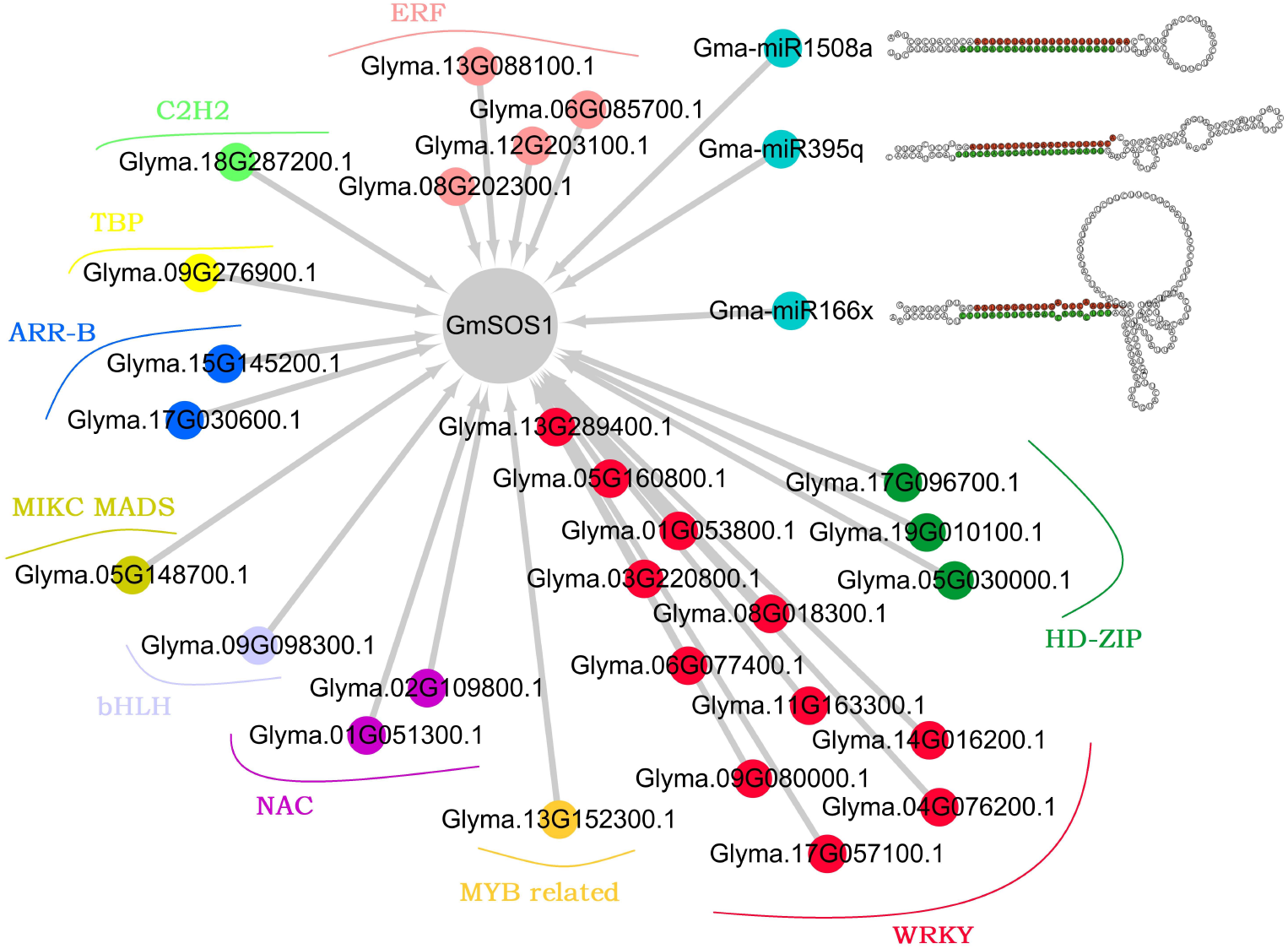
transcription factors and miRNAs that regulate *GmSOS1*.

### 3.7 Conserved motif and phylogenetic analysis

The results of conserved motif analysis showed that the motifs 2∼7 were shared by all SOS1 members, indicating that these motifs are highly conserved in SOS1 (Figure 8). They might be responsible for different biological functions in these SOS1s. Motif2 and motif3 presented in the Na^+^/H^+^ exchanger domain of SOS1s. Motif4、 5、 6 and 7 existed in cyclic nucleotide-binding domain. The conserved C-terminal region harbored more motifs than N-terminal.

**Figure 8.**
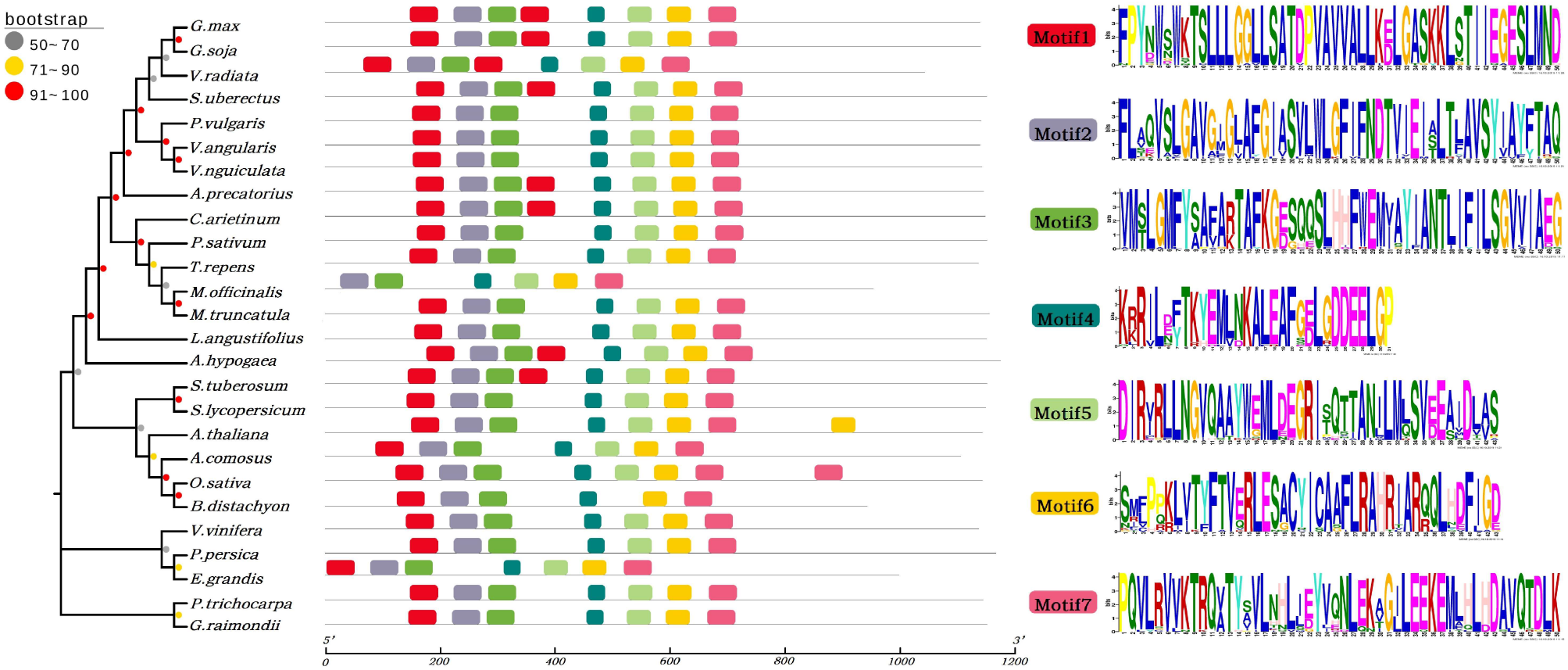
phylogenetic tree and comparison analysis of the conserved motif of SOS1 protein.

Comparative analysis of homologs between *Glycine max* and *Glycine soja* may be helpful for the understanding of homologous protein function in soybean. Our results showed that GmSOS1 and GsSOS1 are clustered into a same branch in the phylogenetic tree (Figure 8), implicating that GmSOS1 protein might be function as GsSOS1. Clearly, protein form *Glycine max* showed more distantly evolutionary relationship with *Gossypium raimondii* and *Populus trichocarpa*, indicating that the differentiation time between them may be the longest, or the selection pressure of SOS1 protein among them is quite different.

## 4. Conclusion

The plasma membrane Na^+^/H^+^ exchanger SOS1 (salt-overly-sensitive 1) is a critical salt tolerance determinant in plants. In this study, a systematic analysis was executed to study the *GmSOS1* gene. Subcellular localization confirmed that soybean *GmSOS1* gene is localized at the plasma membrane, which is consistent with the presumption that GmSOS1 is a plasma membrane Na^+^/H^+^ antiporter. Domain analysis showed that soybean GmSOS1 protein has a Na^+^/H^+^ exchanger domain, a cyclic nucleotide-binding domain, and an auto-inhibitory domain, which are closely related to the function of SOS1 protein to efflux Na^+^. The prediction of the phosphorylation sites combined with the analysis of the transmembrane domain indicates that there are four phosphorylation sites in the soybean GmSOS1 protein, and they are likely to be target sites for protein kinase to regulate GmSOS1 transport activity. The gene co-functional network predicted that *GmCBL4* has the same function as *SOS3, GmCBL10* has the same function as *AtCBL10*, and a new *GmCBL7* that may have similar functions as *SOS3*. The expression of *GmSOS1* gene is spatio-temporal specific, and a variety of transcription factors and miRNAs can regulate its expression. We found that *GmWRKY47* may regulate its transcription by binding to the promoter of *GmSOS1*, which provides an idea for the mechanism study of *GmWRKY47* participating in the response to salt stress. In the analysis of 26 homologous proteins, the highly conserved motif2∼motif7 suggests that the core functional sites of SOS1 might be composed of them. Phylogenetic analysis showed that GmSOS1 is more closely related to GsSOS1. Soybean is an important oil crop, the results of this work are helpful to study the growth and development of soybean under salt stress. Our present study raises several interesting questions that need to be addressed in the future, for example, 1) How *GmSOS1* and *HKT1* work together to regulate intracellular K^+^/Na^+^ balance, 2) How the complex formed by the hydrophilic tail of GmSOS1 and RCD1 plays a role in resisting salt stress, 3) How cyclins are involved in the *GmSOS1* regulatory pathway, 4) What role the predicted transcription factors play in the regulation of *GmSOS1*.

